# Laminar-specific cortical dynamics in human visual and sensorimotor cortices

**DOI:** 10.1101/226274

**Authors:** James J Bonaiuto, Sofie S Meyer, Simon Little, Holly Rossiter, Martina F Callaghan, Fred Dick, Gareth R Barnes, Sven Bestmann

**Author notes:** joint last author (Gareth R Barnes, Sven Bestmann).

## Abstract

Lower frequency, feedback, activity in the alpha and beta range is thought to predominantly originate from infragranular cortical layers, whereas feedforward signals in the gamma range stem largely from supragranular layers. Distinct anatomical and spectral channels may therefore play specialized roles in communication within hierarchical cortical networks; however, empirical evidence for this organization in humans is limited. We leverage high precision MEG to test this proposal, directly and non-invasively, in human participants during visually guided actions. Visual alpha activity mapped onto deep cortical laminae, whereas visual gamma activity predominantly arose from superficial laminae. This laminar-specificity was echoed in sensorimotor beta and gamma activity. Visual gamma activity scaled with task demands in a way compatible with feedforward signaling. For sensorimotor activity, we observed a more complex relationship with feedback and feedforward processes. Distinct frequency channels thus operate in a laminar-specific manner, but with dissociable functional roles across sensory and motor cortices.

## Introduction

The cerebral cortex is hierarchically organized via feedback and feedforward connections that originate predominantly from deep and superficial layers, respectively (Barone et al., 2000; Felleman and Van Essen, 1991; Markov et al., 2013, 2014a, 2014b). Evidence from non-human animal models suggests that information along those pathways is carried via distinct frequency channels: lower frequency (<30Hz) signals predominantly arise from deeper, infragranular layers, whereas higher frequency (>30Hz) signals stem largely from more superficial, supragranular layers (Bollimunta et al., 2008, 2011; Buffalo et al., 2011; Haegens et al., 2015; van Kerkoerle et al., 2014; Maier et al., 2010; Roopun et al., 2006, 2010; Smith et al., 2013; Sotero et al., 2015; Spaak et al., 2012; Sun and Dan, 2009; Xing et al., 2012). These data have inspired general theories of cortical functional organization which ascribe specific computational roles to these pathways (Adams et al., 2013; Arnal and Giraud, 2012; Bastos et al., 2012; Donner and Siegel, 2011; Fries, 2005, 2015; Friston and Kiebel, 2009; Jensen and Mazaheri, 2010; Jensen et al., 2015; Stephan et al., 2017; Wang, 2010). In these proposals, lower frequency activity subserves feedback, top-down communication, and originates in infragranular layers, whereas high-frequency activity is predominantly carried via projections from supragranular layers and conveys feedforward, bottom-up information.

However, evidence for these proposals in humans is largely indirect and focused on visual and auditory areas (Fontolan et al., 2014; Kok et al., 2016; Koopmans et al., 2010; Michalareas et al., 2016; Olman et al., 2012; Scheeringa and Fries, 2017). Whether it is indeed possible to attribute low and high frequency activity in humans to laminar-specific sources, throughout the cortical hierarchy, remains unclear. Here we leverage recent advances in high precision magnetoencephalography (MEG; Meyer et al., 2017a; Troebinger et al., 2014a) to address this issue directly and non-invasively across human visual and sensorimotor cortices.

MEG is a direct measure of neural activity (Baillet, 2017; Hämäläinen et al., 1993), with millisecond temporal precision that allows for delineation of brain activity across distinct frequency bands. Recently developed 3D printed headcast technology gives us more stability in head positioning as well as highly precise models of the underlying cortical anatomy. Together, this allows us to record higher signal-to-noise ratio (SNR) MEG data than previously achievable (Meyer et al., 2017a; Troebinger et al., 2014a). Theoretical and simulation work shows that these gains allow for distinguishing the MEG signal originating from either deep or superficial laminae (Troebinger et al., 2014b), in a time-resolved and spatially localized manner (Bonaiuto et al., 2017). We therefore employed this approach to directly test for the proposed laminar-specificity of distinct frequency channels in human cortex. Such a demonstration would provide important clarification for the proposed mechanism of inter-regional communication in hierarchical cortical networks.

## Results

### Behavioral responses vary with perceptual evidence and cue congruence

We investigated the laminar and spectral specificity of feedforward and feedback signals in visual and sensorimotor cortex with a visually guided action selection task. The task was designed to induce well-studied patterns of low- and high-frequency activity in visual (Busch et al., 2004; Fries et al., 2001; Hari and Salmelin, 1997; Hoogenboom et al., 2006; Mazaheri et al., 2014; Müller et al., 1996; Muthukumaraswamy and Singh, 2013; Sauseng et al., 2005; Thut, 2006; Yamagishi et al., 2005) and sensorimotor cortices (Cheyne et al., 2008; Crone et al., 1998; Donner et al., 2009; Gaetz et al., 2011; Haegens et al., 2011; Huo et al., 2010; de Lange et al., 2013; Pfurtscheller and Neuper, 1997; Pfurtscheller et al., 1996; Tan et al., 2016, 2014; Torrecillos et al., 2015). Participants first viewed a random dot kinetogram (RDK) with coherent motion to the left or the right, which in most trials was congruent to the direction of the following instruction cue indicating the required motor response (an arrow pointing left equated to an instruction to press the left button, and vice versa; **Figure 1A**).

**Figure 1.**
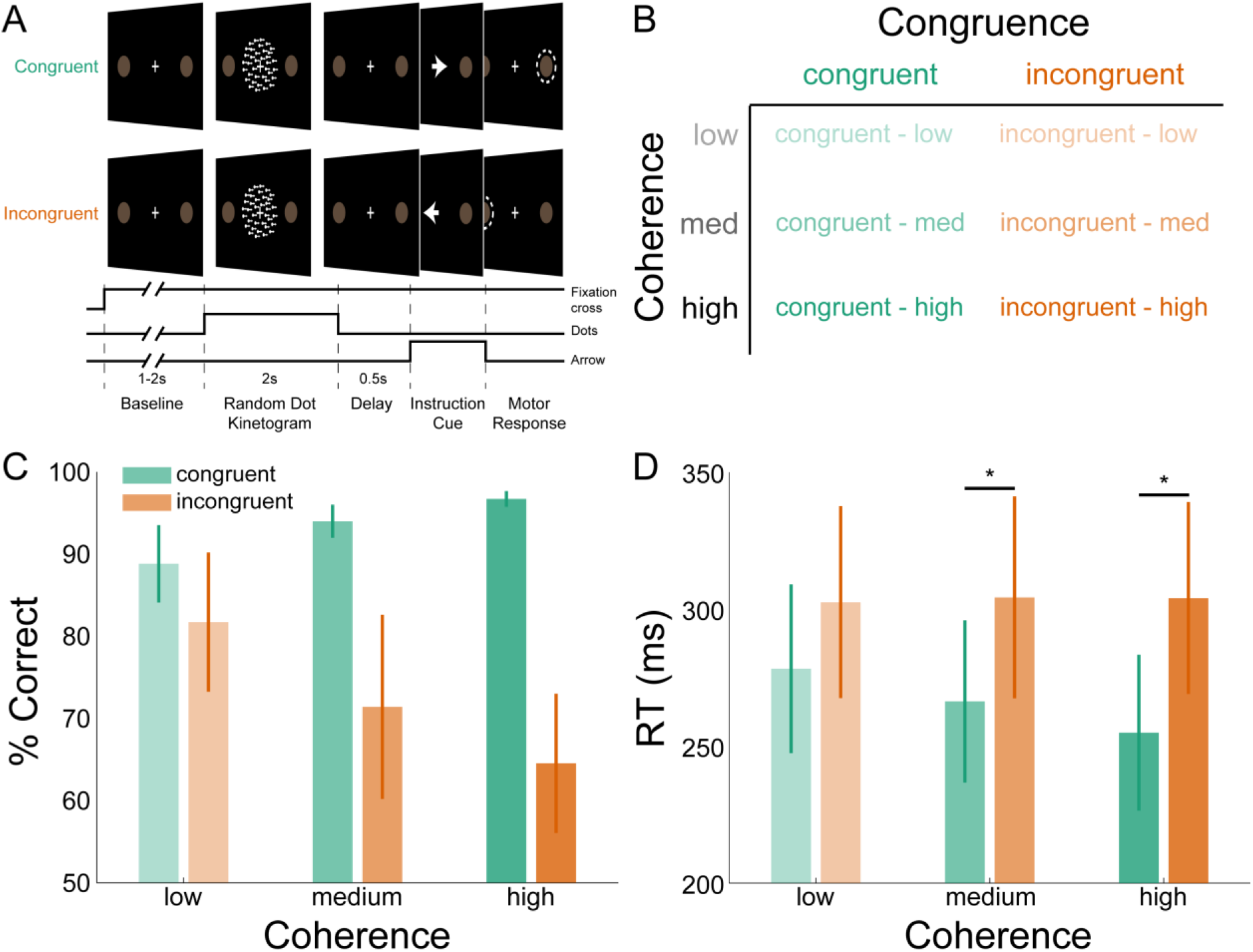
Task structure and participant behavior. A) Each trial consisted of a fixation baseline (1-2s), random dot kinetogram (RDK; 2s), delay (0.5s), and instruction cue intervals, followed by a motor response (left/right button press) in response to the instruction cue (an arrow pointing in the direction of required button press). During congruent trials the coherent motion of the RDK was in the same direction that the arrow pointed in the instruction cue, while in incongruent trials the instruction cue pointed in the opposite direction. B) The task involved a factorial design, with three levels of motion coherence in the RDK and congruent or incongruent instruction cues. Most of the trials (70%) were congruent. C) Mean accuracy over participants during each condition. Error bars denote the standard error. Accuracy increased with increasing coherence in congruent trials, and worsened with increasing coherence in incongruent trials. D) The mean response time (RT) decreased with increasing coherence in congruent trials (* p<0.05).

Participants could therefore accumulate the sensory evidence from the RDK during the 2s it was presented for in order to prepare their response in advance of the instruction cue. However, in incongruent trials the instruction cue was an arrow pointing in the opposite direction from the direction of coherent motion of the RDK, and so the opposite response from the expected one was required. The strength of the motion coherence was varied, modulating the strength of instructed response predictability, and thus we assume, feedforward and feedback activity (**Figure 1B**; Donner et al., 2009; de Lange et al., 2013).

As expected, particpants responded more accurately and more quickly with increasing RDK motion coherence during congruent trials, while behavioral performance worsened with increasing coherence during incongruent trials (**Figure 1**C, D). This was demonstrated by a significant interaction between congruence and coherence for accuracy (F(2,35)=8.201, p=0.004), and RT (*F*(2,35)=7.392, *p*=0.006). Pairwise comparisons (Bonferroni corrected) showed that RTs were faster during congruent trials than incongruent trials at medium (*t*(7)=-3.235, *p*=0.0429) and high coherence levels (*t*(7)=-3.365, *p*=0.036). Participants were thus faster and more accurate when the cued action matched the action they had prepared (congruent trials), and slower and less accurate when these actions were incongruent.

### High SNR MEG recordings using individualized headcasts

Subject-specific headcasts minimize both within-session movement and co-registration error (Meyer et al., 2017a; Troebinger et al., 2014a). This ensures that when MEG data are recorded over separate days, the brain remains in the same location with respect to the MEG sensors. In all participants, within-session movement was <0.2mm in the x and y dimensions and <1.5mm in the z dimension, while co-registration error was <1.5mm in any dimension (estimated by calculating the within-participant standard deviation of the absolute coil locations across recording blocks; **Figure S1**). To assess the between-session homogeneity of our data, we examined topographic maps, event-related fields (ERFs), and time-frequency decompositions. In each, the data were analyzed in three ways: aligned to the onset of the RDK (**Figure 2A**), instruction cue (**Figure 2B**), or button response (**Figure 2C**). The data were acquired during four separate recording sessions, spaced at least a week apart. This analysis revealed that topographic maps and event-related fields from individual MEG sensors and time-frequency spectra from sensor clusters are highly reproducible across different days of recording within an individual participant. Because the headcast approach ensured that participants were in an identical position on repeated days of recording, we were able to obtain very high signal-to-noise (SNR) datasets.

**Figure 2:**
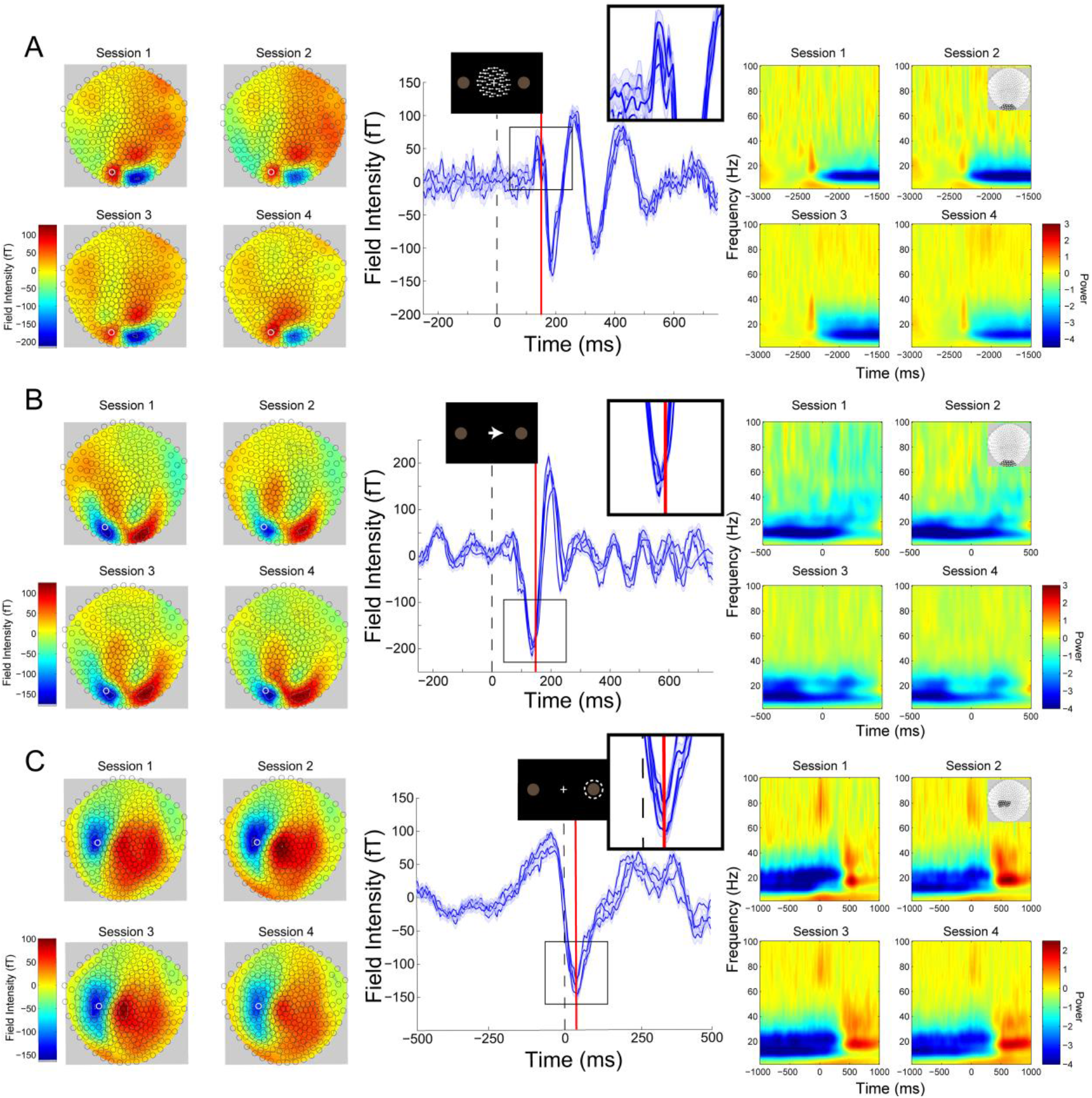
Cross-session reproducibility. Topographic maps (left column), event-related fields (ERFs, middle column), and time-frequency decompositions (right column) aligned to: A) onset of the random dot kinetogram (RDK), B) onset of the instruction cue, or C) the participant’s response (button press). Data shown are for a single representative participant for four sessions on different days (each including three, 15 minute blocks, 180 trials per block). The white circles on the topographic maps denote the sensor from which the ERFs in the middle are recorded. Each blue line in the ERF plots represents a single session (average of 540 trials), with shading representing the standard error (within-session variability) and the red lines showing the time point that the topographic maps are plotted for (150ms for the RDK and instruction cue, 35ms for the response). The insets show a magnified view of the data plotted within the black square. The time-frequency decompositions are baseline corrected (RDK-aligned: [-500, 0ms]; instruction cue-aligned: [-3s, -2.5s]; response-aligned: [-500ms, 0ms relative to the RDK]) and averaged over the sensors shown in the insets.

### Low and high frequency activity localize to different cortical laminae

To address our main question about the laminar specificity of different frequency channels in human cortex, we extracted task-related low- and high-frequency activity from visual and sensorimotor cortices. Attention to visual stimuli is associated with decreases in alpha (Hari and Salmelin, 1997; Mazaheri et al., 2014; Sauseng et al., 2005; Thut, 2006; Yamagishi et al., 2005) and increases in gamma activity in visual cortex (Busch et al., 2004; Fries et al., 2001; Hoogenboom et al., 2006; Müller et al., 1996; Muthukumaraswamy and Singh, 2013). We therefore examined the decrease in alpha (7-13Hz) power following the onset of the RDK, as well as the increase in gamma (60-90Hz) activity following the onset of the RDK and the instruction cue.

Motor responses are associated with a stereotypical pattern of spectral activity in contralateral sensorimotor cortex involving a decrease in beta power during response preparation, followed by a rebound in beta activity. Moreover, a burst of gamma activity typically occurs in contralateral sensorimotor cortex aligned to the movement (Cheyne et al., 2008; Crone et al., 1998; Gaetz et al., 2011; Huo et al., 2010; Pfurtscheller and Neuper, 1997; Pfurtscheller et al., 1996). These two signals are relevant for testing the proposed feedback and feedforward role of low and high frequency activity, respectively, for the following reasons. First, the beta power decrease prior to movement is thought to reflect the removal of inhibition that prevents movement (Engel and Fries, 2010). Moreover, gamma bursts at movement onset arise from motor cortex, are effector-specific, and are thought to reflect the feedback control of discrete movements (Cheyne et al., 2008; Muthukumaraswamy, 2010), and prediction error processing for the purpose of updating motor predictions (Mehrkanoon et al., 2014). The akinetic role of pre-movement beta and the proposed role of movement-related gamma would be difficult to reconcile with the proposed role of these frequency channels in feedback and feedforward control in sensory cortices. This suggests that in sensorimotor cortex, these activity channels may not be organized in the same laminar-specific manner. Alternatively, the same laminar-specific organization may have functional roles that are distinct from the proposed feedback and feedforward communication in sensory cortex. We therefore analyzed the decrease in sensorimotor beta (15-30Hz) power during the RDK and its subsequent rebound following the participant’s response, as well as the response-aligned gamma (60-90Hz) burst.

Localization of activity measured by MEG sensors requires accurate generative forward models which map from cortical source activity to measured sensor data (Baillet, 2017; Hillebrand and Barnes, 2002, 2003; Larson et al., 2014). We constructed a generative model for each participant based on a surface mesh including both their white matter and pial surfaces, representing both deep and superficial cortical laminae, respectively (**Figure 3**, left column). We were thus able to compare the estimated source activity for measured visual and sensorimotor activity on the white matter and pial surface, and infer its laminar origin as deep if the activity is strongest on the white matter surface or superficial if it is strongest on the pial surface. For the purposes of comparison with invasive neural recordings, deep laminae correspond to infragranular cortical layers, and superficial laminae correspond to supragranular layers.

**Figure 3:**
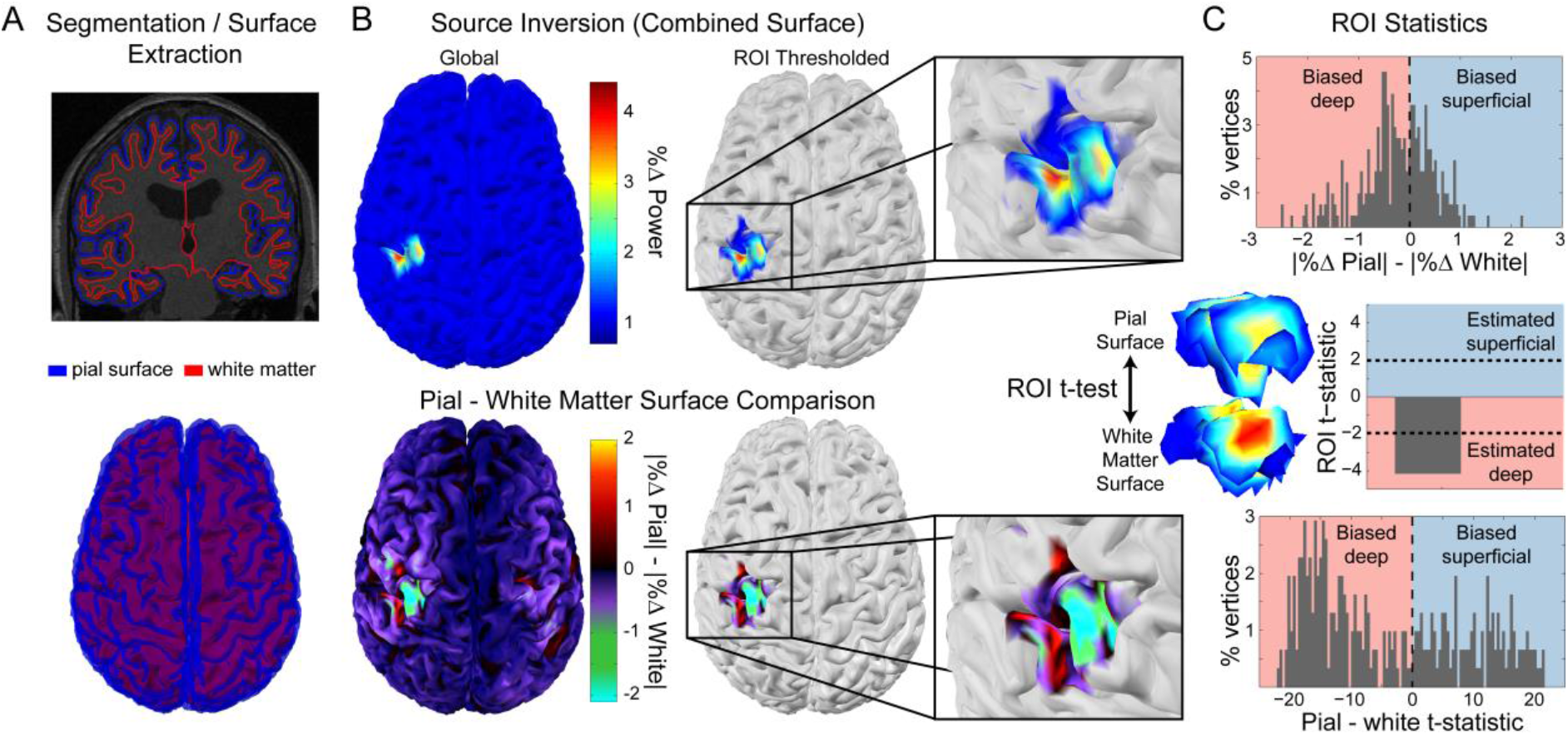
Laminar analysis. Pial and white matter surfaces are extracted from quantitative maps of proton density and T1 times obtained from a multi-parameter mapping MRI protocol (A, top). The model constitutes a generative model combining both surfaces (A, bottom) which is used to perform source inversion using the measured sensor data, resulting in an estimate of the activity at every vertex on each surface (B, top left). The ROI analysis defined a region of interest by comparing the change in power in a particular frequency band during a time window of interest from a baseline time period (B, top right). The ROI includes all vertices in either surface in the 80th percentile as well as corresponding vertices in the other surface. The absolute change in power on each surface was then compared within the ROI (B, bottom; C, top). Pairwise t-tests were performed between corresponding vertices on each surface within the ROI to examine the distribution of t-statistics (C, bottom), as well as on the mean absolute change in power within the ROI on each surface to obtain a single t-statistic which was negative if the greatest change in power occurred on the white matter surface, and positive if it occurred on the pial surface (C, middle).

The veracity of laminar inferences using this analysis is highly dependent on the accuracy of the white matter and pial surface segmentations. Imprecise surface reconstructions from standard 1mm isotropic T1-weighted volumes result in coarse-grained meshes, which do not accurately capture the separation between the two surfaces, and thus do not allow distinctions to be made between deep and superficial laminae (**Figure S2**). We therefore extracted each surface from high-resolution (800μm isotropic) MRI multi-parameter maps (Carey et al., 2017), allowing fine-grained segmentation of the white matter and pial surfaces.

For each low- and high-frequency visual and sensorimotor signal, the laminar analysis first calculated the absolute change in power from a baseline time window on the vertices of each surface, and then compared the power change between surfaces using paired t-tests. The resulting t-statistic was positive when the change in power was greater on the pial surface (superficial), and negative when the change was greater on the white matter surface (deep; **Figure 3**). To get a global measure of laminar specificity, we averaged the change in power over the whole brain (all vertices) within each surface. In order to make spatially localized laminar inferences, we then identified regions of interest (ROIs) in each subject based on the mean frequency-specific change in power from a baseline time window on vertices from either surface (Bonaiuto et al., 2017; **Figure 3**). We further compared two metrics for defining the ROIs: functionally defined (centered on the vertex with the peak mean difference in power), and anatomically-constrained (centered on the vertex with the peak mean power difference within the visual cortex bilaterally, or in the contralateral motor cortex).

### Visual alpha and gamma have distinct laminar specific profiles

Based on *in vivo* laminar recordings in non-human primates (Bollimunta et al., 2008, 2011; Buffalo et al., 2011; Haegens et al., 2015; van Kerkoerle et al., 2014; Maier et al., 2010; Spaak et al., 2012; Sun and Dan, 2009; Xing et al., 2012), we reasoned that changes in alpha activity following the RDK should predominate in infragranular cortical layers. By contrast, changes in gamma activity following the RDK and instruction cue should be strongest in supragranular layers. Source reconstructions of the change in visual alpha activity following the onset of the RDK on the white matter and pial surfaces approximating the proposed laminar origins are shown for an example participant over the whole brain and within the functionally defined ROI in **Figure 4A**. Activity on both surfaces localized to visual cortex bilaterally. When performing paired t-tests comparing corresponding vertices on the pial and white matter surfaces over all trials, the distribution of alpha activity was skewed toward the white matter surface, in line with an infragranular origin. This bias was also observed within the functionally defined ROI. When averaging the change in power either over the whole brain, within a functionally-defined, or an anatomically constrained ROI, the visual alpha activity of most participants was classified as originating from the white matter surface (global: 8/8 participants, functional ROI: 7/8 participants, anatomical ROI: 5/8 participants; **Figure 4A**, right).

**Figure 4:**
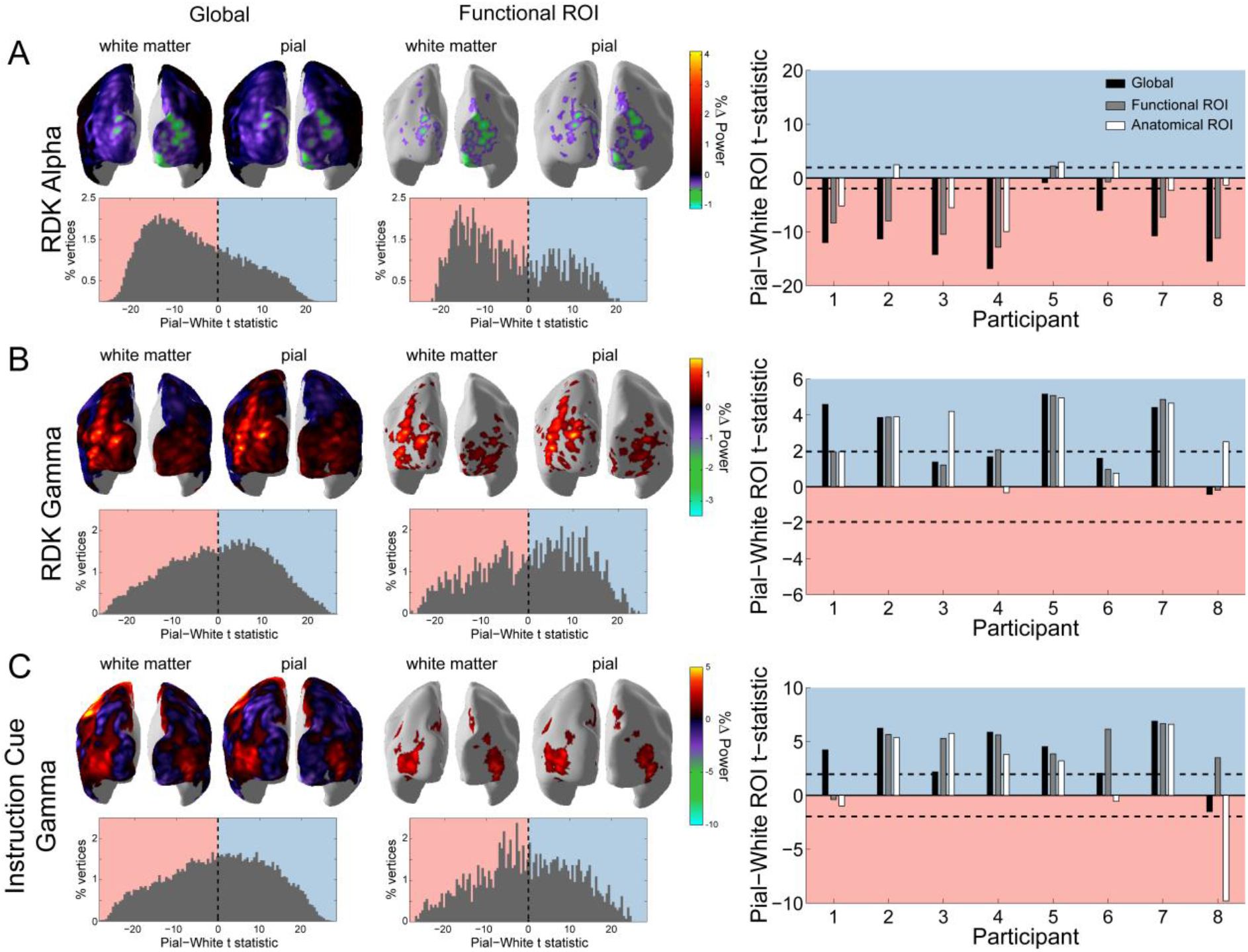
Laminar specificity of visual alpha and gamma. A) Estimated changes in alpha power (7-13Hz) from baseline on the white matter and pial surface following the onset of the random dot kinetogram (RDK), over the whole brain (global) and within a functionally defined region of interest (ROI). Histograms show the distribution of t-statistics comparing the absolute change in power between corresponding pial and white matter surface vertices over the whole brain, or within the ROI. Negative t-statistics indicate a bias toward the white matter surface, and positive t-statistics indicate a pial bias. The bar plots show the t-statistics comparing the absolute change in power between the pial and white matter surfaces averaged within the ROIs, over all participants. T-statistics for the whole brain (black bars), functionally defined (grey bars), and anatomically constrained (white bars) ROIs are shown (red = biased toward the white matter surface, blue = biased pial). Dashed lines indicate the threshold for single subject statistical significance. B) As in A, for gamma (60-90Hz) power following the RDK. C) As in A and B, for gamma (60-90Hz) power following the instruction cue.

Conversely, the increase in visual gamma following the onset of the RDK and instruction cue was strongest on the pial surface (**Figure 4B**, C) as expected. Example source reconstructions on the pial and the white matter surface show activity in the same bilateral areas over visual cortex as visual alpha (**Figure 4B**, C). For visual gamma, the distributions of t-statistics for pairwise vertex comparisons were skewed toward the pial surface, a finding that is compatible with a supragranular origin of high-frequency gamma activity. This was confirmed in subsequent global, functional, and anatomical ROI metrics (RDK gamma, global: 7/8 participants; RDK gamma, functional ROI: 7/8 participants; RDK gamma, anatomical ROI: 7/8 participants; instruction cue gamma, global: 7/8 participants; instruction cue gamma, functional ROI: 7/8 participants; instruction cue gamma, anatomical ROI: 5/8 participants).

We then conducted three control analyses to ascertain the robustness of our findings: shuffling of the position of the sensors, simulation of increased co-registration error, and decreasing effective SNR by using only a random subset of the trials for each participant (see Supplemental Information). Shuffling the position of the sensors destroys any correspondence between the anatomy and the sensor data. Added co-registration error simulates the effect of between-session spatial uncertainty arising from head movement and inaccuracies of the forward model typically experienced without headcasts (Hillebrand and Barnes, 2003, 2011; Medvedovsky et al., 2007; Meyer et al., 2017b; Troebinger et al., 2014b; Uutela et al., 2001). For both control analyses, visual alpha and gamma activity now localized to the pial surface (**Figure S3**, S4), suggesting that the laminar discrimination between visual alpha and gamma in our main analyses would not have been possible were it not for the high-SNR data coupled with the high-precision anatomical models.

The magnitude of the ROI t-statistics for all participants increased with the number of trials used in the analysis, with more trials required for visual gamma signals to reach significance (**Figure S5**). Therefore, the laminar bias exhibited by visual alpha and gamma was unlikely to be driven by a small subset of the trials. One concern was that the effects could be driven by signal power (i.e. higher power signals always localize deeper). Importantly however, regardless of the SNR the shuffled sensor models did not show this behavior within the functionally defined and anatomically constrained ROIs (**Figure S5**).

### Sensorimotor beta and gamma originate from distinct cortical laminae

The above results provide novel support for distinct anatomical pathways through which different frequency channels contribute to intra-areal communication within visual cortex. We next addressed whether this laminar specificity of different frequency channels was common to other portions of cortex, specifically sensorimotor cortex.

Cortical regions vary in terms of thickness (Fischl and Dale, 2000; Jones et al., 2000; Kabani et al., 2001; Lerch and Evans, 2005; MacDonald et al., 2000), as a result of inter-regional variation in cortical folding and laminar morphology (Barbas and Pandya, 1989; Hilgetag and Barbas, 2006; Matelli et al., 1991; Rajkowska and Goldman-Rakic, 1995). Moreover, the distinction between feedback and feedforward cortical processing channels may be less clear for motor cortex, which is agranular (missing layer IV) and projects directly to the spinal cord. Supporting this argument, motor gamma bursts are closely tied to movement onset, and thought to reflect the execution, or feedback control, of movement (Cheyne and Ferrari, 2013; Cheyne et al., 2008).

While frequency-specific activity thus occurs throughout cortex, the laminar distribution of different frequency channels may differ across different levels in the cortical hierarchy. Because MEG is only sensitive to the synchronous activity of large populations of pyramidal cells, it is likely that different laminar microcircuits could give rise to the same measurable MEG signals (Cohen, 2017). Alternatively, if the layer specificity of low and high frequency activity is a general organizing principle of cortex, one would expect the pre-movement beta decrease and post-movement rebound to originate from infragranular cortical layers, and the movement-related gamma increase to be strongest in supragranular layers. Moreover, the ability of MEG to accurately segregate deep from superficial laminar source activity may vary throughout cortex, a possibility we have previously explored in simulation (Bonaiuto et al., 2017).

To explore this possibility empirically, we analyzed two task-related modulations of sensorimotor beta activity: the decrease in beta power following the onset of the RDK, just prior to the motor response, and the post-movement beta rebound (Cassim et al., 2001; Jurkiewicz et al., 2006; Parkes et al., 2006; Pfurtscheller et al., 1996; Salmelin et al., 1995). Both signals localized to the left sensorimotor cortex (contralateral to the hand used to indicate the response; **Figure 5A**, B), and both signals were strongest on the white matter surface, as evidenced by the white matter skews in the global and functional ROI t-statistics (**Figure 5**). This laminar pattern with both the beta decrease and rebound classified as originating from the white matter surface held for all but one participant. This general finding is of relevance as it addresses concerns that the high SNR of beta activity trivially leads to its attribution to the deeper cortical surface. Here, the two epochs of beta activity were characterized by power decreases and increases, respectively, meaning that SNR alone cannot explain the laminar localization of beta activity.

**Figure 5:**
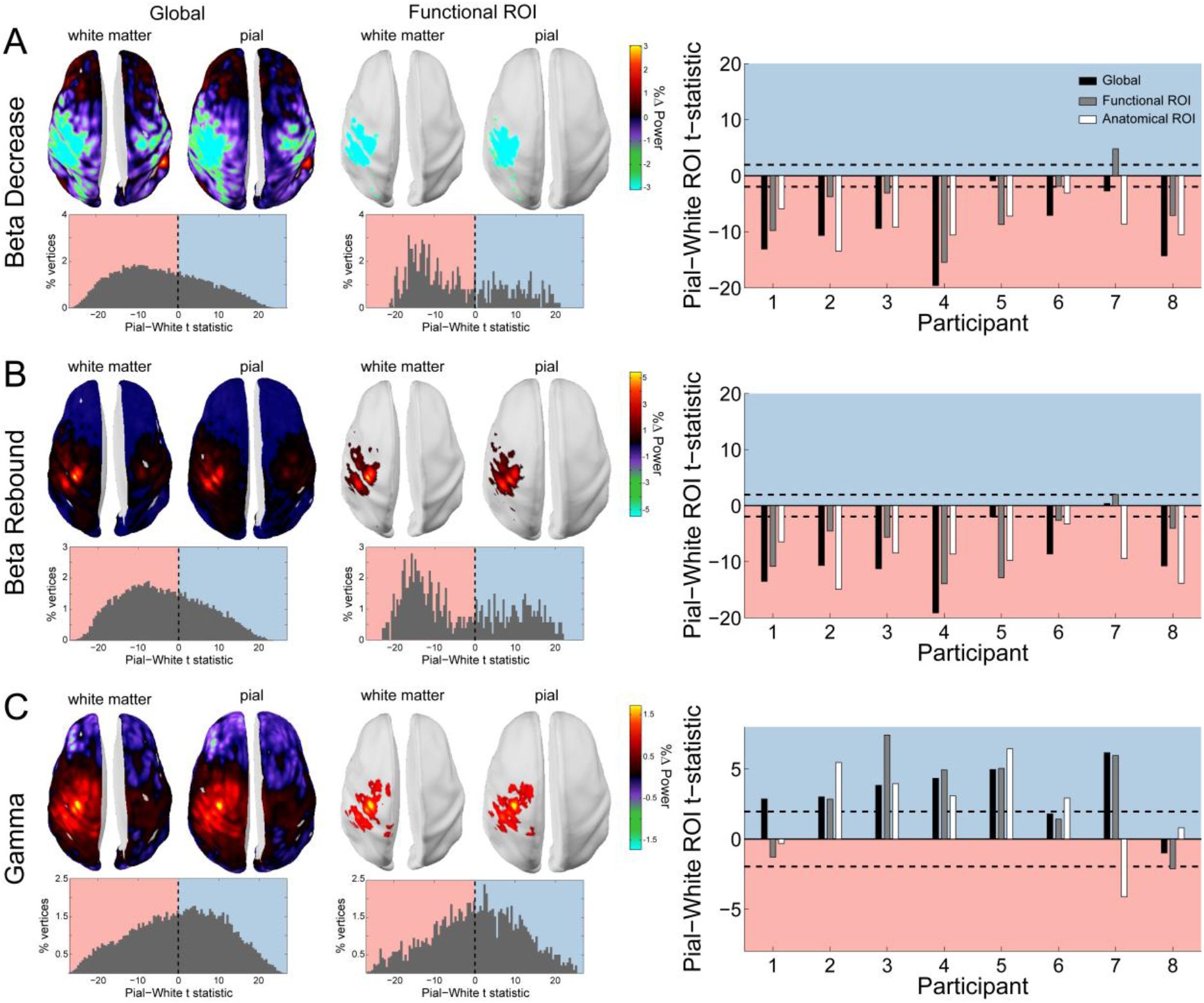
Laminar specificity of sensorimotor beta and gamma. As in figure 4, for A) the beta (15-30Hz) decrease prior to the response, B) beta (15-30Hz) rebound following the response, and C) gamma (60-90Hz) power change from baseline during the response. In the histograms and bar plots, positive and negative values indicate a bias towards the superficial and deeper cortical layers, respectively. The dashed lines indicate single subject level significance thresholds. The black, grey, and white bars indicate statistics based on regions of interest comprising the whole brain, functional and anatomically-constrained ROIs, respectively.

The burst of gamma aligned with the onset of the movement localized to the same patch of left sensorimotor cortex (**Figure 5C**), and in the example participant shown in **Figure 5** and for most participants, was strongest on the pial surface (global: 7/8 participants; function ROI: 6/8 participants; anatomical ROI: 6/8 participants).

Next we repeated our control analysis on the sensorimotor data, which mirrored those of visual alpha and gamma. Sensor shuffling, as well as the addition of co-registration error, resulted in sensorimotor beta and gamma localizing to the pial surface (**Figure S3, S4**), and the ROI t-statistics increased in magnitude with the number of trials used in the analysis, with more trials required for sensorimotor gamma signals to pass the significance threshold (**Figure S5**). Again, importantly, the gamma superficial bias within the functionally defined and anatomically constrained ROIs did not increase with SNR for the shuffled sensor data, meaning that the superficial localization of gamma was not driven by low SNR (**Figure S5**).

### Superficial visual gamma scales with cue congruence

Next, we asked whether the observed low and high-frequency laminar-specific activity in visual and sensorimotor cortex dynamically varied with task demands in line with proposals about their role in feedback and feedforward message passing (Adams et al., 2013; Arnal and Giraud, 2012; Bastos et al., 2012; Donner and Siegel, 2011; Fries, 2005, 2015; Friston and Kiebel, 2009; Jensen and Mazaheri, 2010; Jensen et al., 2015; Stephan et al., 2017; Wang, 2010). This would provide additional indirect support for the idea that communication in hierarchical cortical networks is organized through distinct frequency channels along distinct anatomical pathways, to orchestrate top-down and bottom-up control.

In our task, the direction of the instruction cue was congruent with the motion coherence direction in the RDK during most trials (70%). As such, if the direction of motion coherence is to the left, the instruction cue will most likely be a leftward arrow. Gamma activity increases in sensory areas during presentation of unexpected stimuli (Arnal et al., 2011; Gurtubay et al., 2001; Todorovic et al., 2011), and therefore we expected visual gamma activity in supragranular layers to be greater following incongruent instruction cues than after congruent cues. Indeed, the increase in visual gamma on the pial surface following the onset of the instruction cue was greater in incongruent compared to congruent trials (W(8)=0, p=0.008; 8/8 participants; incongruent-congruent M=1.64%, SD=2.34%; **Figure 6**).

**Figure 6:**
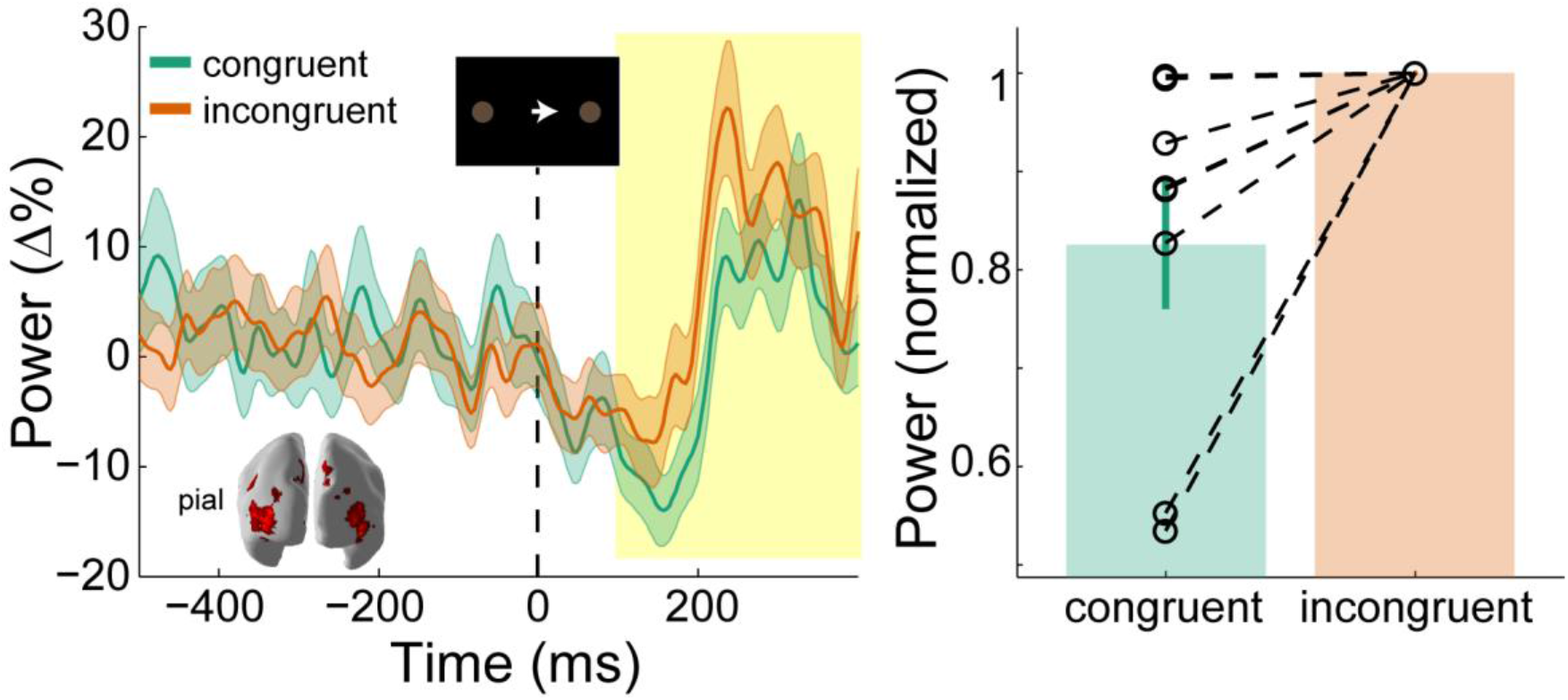
Visual gamma activity modulation by task condition. Visual gamma activity following the onset of the instruction stimulus within the functionally defined ROI of an example participant (left), and averaged within the time window represented by the shaded yellow rectangle for all participants (right). Each dashed line on the right shows the change in normalized values for the different conditions for each participant. The bar height represents the mean normalized change in gamma power, and the error bars denote the standard error. Visual gamma activity is stronger following the onset of the instruction cue when it is incongruent to the direction of the coherent motion in the random dot kinetogram (RDK).

### Deep sensorimotor beta scales with RDK motion coherence and cue congruence

Changes in sensorimotor beta power during response preparation predict forthcoming motor responses (Donner et al., 2009; Haegens et al., 2011; de Lange et al., 2013), whereas the magnitude of sensorimotor beta rebound is attenuated by movement errors (Tan et al., 2014, 2016; Torrecillos et al., 2015). We therefore predicted that, in infragranular layers, the decrease in sensorimotor beta would scale with the motion coherence of the RDK, and the magnitude of the beta rebound would be decreased during incongruent trials when the prepared movement has to be changed in order to make a correct response.

The behavioral results presented thus far suggest that participants accumulated perceptual evidence from the RDK in order to prepare their response prior to the onset of the instruction cue. This preparation was accompanied by a reduction in beta power in the sensorimotor cortex contralateral to the hand used to indicate the response (**Figure 5A**). This beta decrease began from the onset of the RDK and was more pronounced with increasing coherence, demonstrating a significant effect of coherence on the white matter surface (**Figure 7A**; *X*^2^(2)=9.75, p=0.008), with beta during high coherence trials significantly lower than during low coherence trials (8/8 participants; *t*(7)=-3.496, *p*=0.033; low-high M=2.42%, SD=1.96%). Following the response, there was an increase in beta in contralateral sensorimotor cortex (beta rebound) which was greater in congruent, compared to incongruent trials on the white matter surface (**Figure 7B**; W(8)=34, *p*=0.023; 7/8 participants, congruent-incongruent M=5.13%, SD=5.19%). In other words, the beta rebound was greatest when the cued response matched the prepared response.

**Figure 7:**
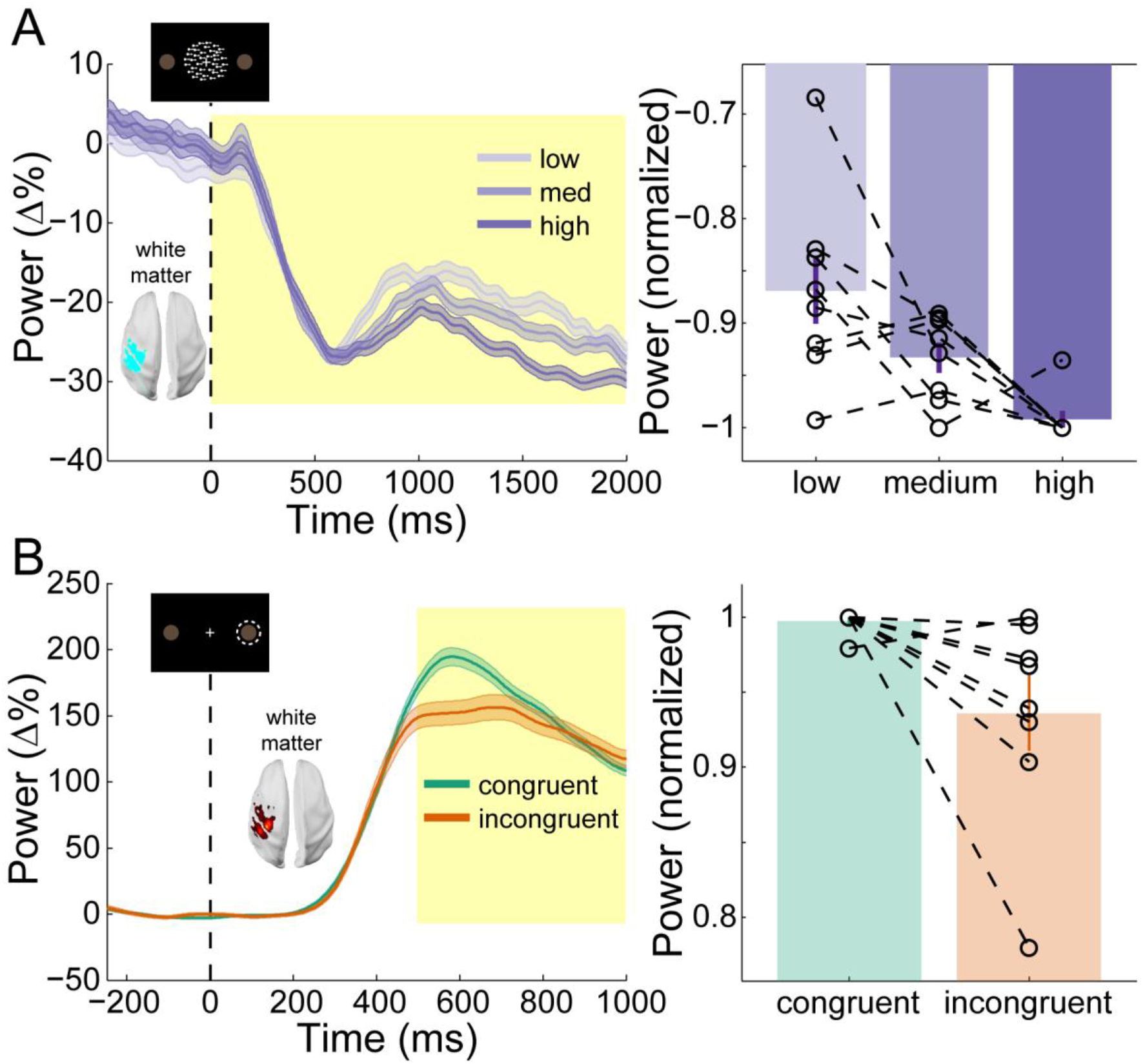
Sensorimotor beta activity modulated by task condition. A) Beta decrease following the onset of the random dot kinetogram (RDK) within the functionally defined ROI of an example participant over the duration of the RDK (left), and averaged over this duration for all participants (right). The bar height represents the mean normalized change in gamma power, and the error bars denote the standard error. The beta decease becomes more pronounced with increasing coherence. B) As in A, for beta rebound following the response and averaged within the time window shown by the shaded yellow rectangle. Beta rebound is stronger following responses in congruent trials.

## Discussion

We have demonstrated that low and high frequency channels localize predominantly to deep and superficial laminae, respectively, in human visual and sensorimotor cortex. These channels play distinct roles in feedback and feedforward processing during visually guided action selection, with high frequency visual activity enhanced by a mismatch between feedforward and feedback signals, and low frequency sensorimotor activity modulated by a combination of feedforward and feedback influences during different task epochs. Through the use of novel MEG headcast technology (Meyer et al., 2017a; Troebinger et al., 2014a) and spatially and temporally resolved laminar analyses (Bonaiuto et al., 2017; Troebinger et al., 2014b), we provide novel evidence for the layer- and frequency-specific accounts of hierarchical cortical organization in humans.

### Low and high frequency channels localize to deep and superficial cortical laminae across visual and sensorimotor cortex

We found that low frequency activity (alpha, 7-13Hz; and beta, 15-30Hz) predominately originated from deep cortical laminae, and high frequency activity (gamma, 60-90Hz) from superficial laminae in both visual and sensorimotor cortex. Our analysis included two built-in controls. Firstly, visually induced gamma after both the RDK and the instruction cue localized superficially, reinforcing the proposal that visual gamma generally predominates from superficial laminae. Secondly, both a decrease and increase in sensorimotor beta power localized to deep laminae, meaning that the laminar analysis was not simply biased toward deep sources for high power signals. Moreover, this laminar specificity was abolished by shuffling the sensors (**Figure S3**) or introducing co-registration error (**Figure S4**), underlining the need for spatially precise anatomical data and MEG recordings. Finally, the laminar bias of both low and high frequency signals increased monotonically as the number of trials included in the analysis increased, but not when the sensors were shuffled (**Figure S5**).

The localization of alpha activity to predominately deep laminae of visual cortex is in line with evidence from depth electrode recordings in visual areas of the non-human primate brain (Buffalo et al., 2011; van Kerkoerle et al., 2014; Maier et al., 2010; Smith et al., 2013; Spaak et al., 2012; Xing et al., 2012). Several studies who have found alpha generators in both infra- and supragranular layers in primary sensory areas (Bollimunta et al., 2008, 2011; Haegens et al., 2015), and it has been suggested that this discrepancy is due to a contamination of infragranular layer LFP signals by volume conduction from strong alpha generators in supragranular layers (Haegens et al., 2015; Halgren et al., 2017). This is unlikely to apply to the results presented here as this type of laminar MEG analysis is biased toward superficial laminae when SNR is low (**Figure S3, S4**; Bonaiuto et al., 2017). However, this analysis can only determine the laminar origin of the strongest activity when it occurs simultaneously at multiple depths (Bonaiuto et al., 2017), which is consistent with the fact that infragranular cortical layers contain the primary local pacemaking alpha generators (Bollimunta et al., 2008, 2011).

We found that gamma activity was strongest in superficial cortical laminae, which was expected given that gamma activity has been found to predominantly occur in supragranular layers in visual cortex (Buffalo et al., 2011; van Kerkoerle et al., 2014; Smith et al., 2013; Spaak et al., 2012; Xing et al., 2012), but see (Nandy et al., 2017). The mechanisms underlying the generation of gamma activity are diverse across the cortex (Buzsáki and Wang, 2012), but commonly involve reciprocal connections between pyramidal cells and interneurons, or between interneurons (Tiesinga and Sejnowski, 2009; Whittington et al., 2011). The local recurrent connections necessary for such reciprocal interactions are most numerous in supragranular layers (Buzsáki and Wang, 2012), as are fast-spiking interneurons which play a critical role in generating gamma activity (Cardin et al., 2009; Carlén et al., 2012; Sohal et al., 2009).

It is widely hypothesized that the laminar segregation of frequency specific channels is a common organizing principle across the cortical hierarchy (Arnal and Giraud, 2012; Bastos et al., 2012; Fries, 2015; Wang, 2010). However, most evidence for this claim comes from depth electrode recordings in primary sensory areas, with the vast majority in visual cortical regions (Buffalo et al., 2011; van Kerkoerle et al., 2014; Smith et al., 2013; Spaak et al., 2012; Xing et al., 2012). While *in vivo* laminar data from primate sensorimotor cortex are lacking, *in vitro* recordings from somatosensory and motor cortices demonstrate that beta activity is generated in neural circuits dominated by infragranular layer V pyramidal cells (Roopun et al., 2006, 2010; Yamawaki et al., 2008). By contrast, gamma activity is thought to arise from supragranular layers II/III of mouse somatosensory cortex (Cardin et al., 2009; Carlén et al., 2012). The results presented here support generalized theories of laminar organization across cortex, and are the first to non-invasively provide evidence for the laminar origin of movement-related sensorimotor activity.

### High frequency activity in visual cortex is enhanced by mismatches in feedforward and feedback signals

We found that visual gamma was enhanced following the presentation of the instruction cue in incongruent compared to congruent trials. This was in agreement with our predictions, based on the fact that supragranular layer gamma activity is implicated in feedforward processing (van Kerkoerle et al., 2014). In our task, the direction of coherent motion in the RDK was congruent with the direction of the following instruction cue in most trials. Participants could therefore form a sensory expectation of the direction of the forthcoming instruction cue, which was violated in incongruent trials. The enhancement of visual gamma following incongruent cues is therefore consistent with the gamma activity increase observed in sensory areas during perceptual expectation violations (Arnal et al., 2011; Gurtubay et al., 2001; Todorovic et al., 2011) as well as layer-specific synaptic currents in supragranular cortical layers during performance error processing (Sajad et al., 2017).

### Low frequency activity in sensorimotor cortex reflects a combination of feedforward and feedback processes

There are numerous theories for the computational role of beta activity in motor systems. Decreases in beta power prior to the onset of a movement predict the selected action (Donner et al., 2009; Haegens et al., 2011; de Lange et al., 2013), whereas the beta rebound following a movement is attenuated by error monitoring processes (Tan et al., 2014, 2016; Torrecillos et al., 2015). Our results unify both of these accounts, showing that the level of beta decrease prior to a movement is modulated by the accumulation of sensory evidence predicting the cued movement, while the beta rebound is diminished when the prepared action must be suppressed in order to correctly perform the cued action. This suggests that in the sensorimotor system, low frequency activity can reflect both bottom-up and top-down processes depending on the task epoch. This may occur via bottom-up, feedforward projections from intraparietal regions to motor regions (Hanks et al., 2006; Kayser et al., 2010; Platt and Glimcher, 1999; Tosoni et al., 2008) or top-down, feedback projections from the dorsolateral prefrontal cortex (Curtis and Lee, 2010; Georgiev et al., 2016; Heekeren et al., 2006, 2004; Hussar and Pasternak, 2013). The dissociation between bottom-up and top-down influences during different task epochs could indicate that the decrease in beta and the following rebound are the result of functionally distinct processes.

### Future directions

Our ROI-based comparison of deep and superficial laminae can only determine the origin of the strongest source of activity, which does not imply that activity within a frequency band is exclusively confined to either deep or superficial sources within the same patch of cortex (Bollimunta et al., 2011; Haegens et al., 2015; Maier et al., 2010; Smith et al., 2013; Spaak et al., 2012; Xing et al., 2012). We should also note that in all of our control studies, in which we discard spatial information, a bias towards the superficial (pial) cortical surface was present. However, this bias does not increase with SNR for high frequency activity with poor anatomical models, mirroring the results of simulations showing that this type of laminar analysis is biased superficially at low SNR levels, but that the metrics are not statistically significant at these levels (Bonaiuto et al., 2017). Moreover, we used white matter and pial surface meshes to represent deep and superficial cortical laminae, respectively, and therefore our analysis is insensitive to granular sources. Recent studies have shown that beta, and perhaps gamma, activity is generated by stereotyped patterns of proximal and distal inputs to infragranular and supragranular pyramidal cells (Jones, 2016; Lee and Jones, 2013; Sherman et al., 2016). Future extensions to our laminar analysis could use a sliding time window in order determine the time course of laminar activity. MEG is a global measure of neural activity, and therefore uniquely situated to test large scale computational models of laminar and frequency-specific interactions (Lee et al., 2013; Mejias et al., 2016; Pinotsis et al., 2017; Wang et al., 2013), as well as the possibility that other cortical areas are organized along different principles; for example, in inferior temporal cortex the primary local pacemaking alpha generators are in supragranular layers (Bollimunta et al., 2008). Finally, in the task used here, participants were told that the direction of coherent motion in the RDK predicts the forthcoming instruction cue. Further research will determine how predictive cues are learned implicitly, and how this process shapes beta and gamma activity in visual and sensorimotor areas.

## Experimental Procedures

### Behavioral Task

Eight neurologically healthy volunteers participated in the experiment (6 male, aged 28.5±8.52 years). The study protocol was in full accordance with the Declaration of Helsinki, and all participants gave written informed consent after being fully informed about the purpose of the study. The study protocol, participant information, and form of consent, were approved by the local ethics committee (reference number 5833/001). Participants completed a visually guided action decision making task in which they responded to visual stimuli projected on a screen by pressing one of two buttons on a button box using the index and middle finger of their right hand. On each trial, participants were required to fixate on a small white cross in the center of a screen. After a baseline period randomly varied between 1s and 2s, a random dot kinetogram (RDK) was displayed for 2s with coherent motion either to the left or to the right (**Figure 1A**). Following a 500ms delay, an instruction cue appeared, consisting of an arrow pointing either to the left or the right, and participants were instructed to press the corresponding button (left or right) as quickly and as accurately as possible. Trials ended once a response had been made or after a maximum of 1s if no response was made.

The task had a factorial design with congruence (whether or not the direction of the instruction cue matched that of the coherent motion in the RDK) and coherence (the percentage of coherently moving dots in the RDK) as factors (**Figure 1B**). Participants were instructed that in most of the trials (70%), the direction of coherent motion in the RDK was congruent to the direction of the instruction cue. Participants could therefore reduce their mean response time (RT) by preparing to press the button corresponding to the direction of the coherent motion. The RDK consisted of a 10°×10° square aperture centered on the fixation point with 100, 0.3° diameter dots, each moving at 4°/s. The levels were individually set for each participant by using an adaptive staircase procedure (QUEST; Watson and Pelli, 1983) to determine the motion coherence at which they achieved 82% accuracy in a block of 40 trials at the beginning of each session, in which they had to simply respond with the left or right button to leftwards or rightwards motion coherence. The resulting level of coherence was then used as medium, and 50% and 150% of it as low and high, respectively.

Each block contained 126 congruent trials, and 54 incongruent trials, and 60 trials for each coherence level with half containing coherent leftward motion, and half rightward (180 trials total). All trials were randomly ordered. Participants completed 3 blocks per session, and 1-5 sessions on different days, resulting in 540-2700 trials per participant (M=1822.5, SD=813.21). The behavioral task was implemented in MATLAB (The MathWorks, Inc., Natick, MA) using the Cogent 2000 toolbox (http://www.vislab.ucl.ac.uk/cogent.php).

### MRI Acquisition

Prior to MEG sessions, participants underwent two of MRI scanning protocols during the same visit: one for the scan required to generate the scalp image for the headcast, and a second for MEG source localization. Structural MRI data were acquired using a 3T Magnetom TIM Trio MRI scanner (Siemens Healthcare, Erlangen, Germany). During the scan, the participant lay in the supine position with their head inside a 12-channel coil. Acquisition time was 3 min 42 s, plus a 45 s localizer sequence.

The first protocol was used to generate an accurate image of the scalp for headcast construction (Meyer et al., 2017a). This used a T1-weighted 3D spoiled fast low angle shot (FLASH) sequence with the following acquisition parameters: 1mm isotropic image resolution, field-of view set to 256, 256, and 192 mm along the phase (anterior-posterior, A–P), read (head-foot, H–F), and partition (right-left, R–L) directions, respectively. The repetition time was 7.96ms and the excitation flip angle was 12°. After each excitation, a single echo was acquired to yield a single anatomical image. A high readout bandwidth (425Hz/pixel) was used to preserve brain morphology and no significant geometric distortions were observed in the images. Acquisition time was 3 min 42s, a sufficiently short time to minimize sensitivity to head motion and any resultant distortion. Care was also taken to prevent distortions in the image due to skin displacement on the face, head, or neck, as any such errors could compromise the fit of the headcast. Accordingly, a more spacious 12 channel head coil was used for signal reception without using either padding or headphones.

The second protocol was a quantitative multiple parameter mapping (MPM) protocol, consisting of 3 differentially-weighted, RF and gradient spoiled, multi-echo 3D FLASH acquisitions acquired with whole-brain coverage at 800μm isotropic resolution. Additional calibration data were also acquired as part of this protocol to correct for inhomogeneities in the RF transmit field (Callaghan et al., 2015; Lutti et al., 2010, 2012). For this protocol, data were acquired with a 32-channel head coil to increase SNR.

The FLASH acquisitions had predominantly proton density (PD), T1 or magnetization transfer (MT) weighting. The flip angle was 6° for the PD- and MT-weighted volumes and 21° for the T1 weighted acquisition. MT-weighting was achieved through the application of a Gaussian RF pulse 2 kHz off resonance with 4 ms duration and a nominal flip angle of 220° prior to each excitation. The field of view was set to 224, 256, and 179 mm along the phase (A–P), read (H–F), and partition (R–L) directions, respectively. Gradient echoes were acquired with alternating readout gradient polarity at eight equidistant echo times ranging from 2.34 to 18.44 ms in steps of 2.30 ms using a readout bandwidth of 488 Hz/pixel. Only six echoes were acquired for the MT-weighted acquisition in order to maintain a repetition time (TR) of 25 ms for all FLASH volumes. To accelerate the data acquisition and maintain a feasible scan time, partially parallel imaging using the GRAPPA algorithm (Griswold et al., 2002) was employed with a speed-up factor of 2 and forty integrated reference lines in each phase-encoded direction (A-P and R-L).

To maximize the accuracy of the measurements, inhomogeneity in the transmit field was mapped by acquiring spin echoes and stimulated echoes across a range of nominal flip angles following the approach described in Lutti et al. (2010), including correcting for geometric distortions of the EPI data due to B0 field inhomogeneity. Total acquisition time for all MRI scans was less than 30 min.

Quantitative maps of proton density (PD), longitudinal relaxation rate (R1 = 1/T1), magnetization transfer saturation (MT) and effective transverse relaxation rate (R2* = 1/T2*) were subsequently calculated according to the procedure described in Weiskopf et al. (2013). Each quantitative map was co-registered to the scan used to design the headcast, using the T1 weighted map. The resulting maps were used to extract cortical surface meshes using FreeSurfer (see below).

### Headcast Construction

From an MRI-extracted image of the skull, a headcast that fit between the participant’s scalp and the MEG dewar was constructed (Meyer et al., 2017a; Troebinger et al., 2014a). Scalp surfaces were first extracted from the T1-weighted MRI scans acquired in the first MRI protocol using standard SPM12 procedures (http://www.fil.ion.ucl.ac.uk/spm/). Next, this tessellated surface was converted into the standard template library (STL) format, commonly used for 3D printing. Importantly, this conversion imposed only a rigid body transformation, meaning that it was easily reverse-transformable at any point in space back into native MRI space. Accordingly, when the fiducial locations were optimized and specified in STL space as coil-shaped protrusions on the scalp, their exact locations could be retrieved and employed for co-registration. Next, the headcast design was optimized by accounting for factors such as headcast coverage in front of the ears, or angle of the bridge of the nose. To specify the shape of the fiducial coils, a single coil was 3D scanned and three virtual copies of it were placed at the approximate nasion, left peri-auricular (LPA), and right peri-auricular (RPA) sites, with the constraint that coil placements had to have the coil-body and wire flush against the scalp, in order to prevent movement of the coil when the headcast was worn. The virtual 3D model was placed inside a virtual version of the scanner dewar such that the distance to the sensors was minimized (by placing the head as far up within the dewar as possible) while ensuring that vision was not obstructed. Next, the head-model (plus spacing elements and coil protrusions) was printed using a Zcorp 3D printer (Zprinter 510) with 600 x 540 dots per inch resolution. The 3D printed head model was then placed inside the manufacturer-provided replica of the dewar and liquid resin was poured in between the surfaces to fill the negative space, resulting in the subject-specific headcast. The fiducial coil protrusions in the 3D model now become indentations in the resulting headcast, in which the fiducial coils can sit during scanning. The anatomical landmarks used for determining the spatial relationship between the brain and MEG sensors are thus in the same location for repeated scans, allowing data from multiple sessions to be combined (Meyer et al., 2017a).

### FreeSurfer Surface Extraction

FreeSurfer (v5.3.0; Fischl, 2012) was used to extract cortical surfaces from the multi-parameter maps. Use of multi-parameter maps as input to FreeSurfer can lead to localized tissue segmentation failures due to boundaries between the pial surface, dura matter and CSF showing different contrast compared to that assumed within FreeSurfer algorithms (Lutti et al., 2014). Therefore, an in-house FreeSurfer surface reconstruction procedure was used to overcome these issues, using the PD and T1 maps as inputs. Detailed methods for cortical surface reconstruction can be found in Carey et al. (Carey et al., 2017). This process yields surface extractions for the pial surface (the most superficial layer of the cortex adjacent to the cerebro-spinal fluid, CSF), and the white/grey matter boundary (the deepest cortical layer). Each of these surfaces is downsampled by a factor of 10, resulting in two meshes comprising about 30,000 vertices each (M=30,094.75, SD=2,665.45 over participants). For the purpose of this paper, we will use these two surfaces to represent deep (white/grey interface) and superficial (grey-CSF interface) cortical models.

### MEG Acquisition

MEG recordings were made using a 275-channel Canadian Thin Films (CTF) MEG system with superconducting quantum interference device (SQUID)-based axial gradiometers (VSM MedTech, Vancouver, Canada) in a magnetically shielded room. The data collected were digitized continuously at a sampling rate of 1200 Hz. A projector displayed the visual stimuli on a screen (~8m from the participant), and participants made responses with a button box.

### Behavioral Analyses

Participant responses were classified as correct when the button pressed matched the direction of the instruction cue, and incorrect otherwise. The response time (RT) was measured as the time of button press relative to the onset of the instruction cue. Both measures were analyzed using repeated measures ANOVAs with congruence (congruent or incongruent) and coherence (low, medium, and high) as factors. Pairwise follow-up tests were performed between congruence levels at each coherence level, Bonferroni corrected.

### MEG Preprocessing

All MEG data preprocessing and analyses were performed using SPM12 (http://www.fil.ion.ucl.ac.uk/spm/) using Matlab R2014a and are available at http://github.com/jbonaiuto/meg-laminar. The data were filtered (5th order Butterworth bandpass filter: 2-100 Hz) and downsampled to 250 Hz. Eye-blink artifacts were removed using multiple source eye correction (Berg and Scherg, 1994). Trials were then epoched from 1s before RDK onset to 1.5s after instruction cue onset, and from 2s before the participant’s response to 2s after. Blocks within each session were merged, and trials whose variance exceeded 2.5 standard deviations from the mean were excluded from analysis.

### Source reconstruction

Source inversion was performed using the empirical Bayesian beamformer (EBB; Belardinelli et al., 2012; López et al., 2014) within SPM. The sensor data were first reduced into 180 orthogonal spatial (lead field) modes and 16 temporal modes. The empirical Bayes optimization rests upon estimating hyper-parameters which express the relative contribution of source and sensor level covariance priors to the data (López et al., 2014). We assumed the sensor level covariance to be an identity matrix, with a single source level prior estimated from the data. The source level prior was based on the beamformer power estimate across a two-layer manifold comprised of pial and white cortical surfaces with source orientations defined as normal to the cortical surface. There were therefore only two hyper-parameters to estimate – defining the relative contribution of the source and sensor level covariance components to the data. We used the Nolte single shell head model as implemented in SPM (Nolte, 2003).

### Analyses for Laminar Discrimination

The laminar analysis reconstructed the data onto a mesh combining the pial and white matter surfaces, thus providing an estimate of source activity on both surfaces (**Figure 3**). We analyzed six different visual and sensorimotor signals at different frequencies and time windows of interest (WOIs): RDK-aligned visual alpha (7-13Hz; WOI=[0s, 2s]; baseline WOI=[-1s, -.5s]), RDK-aligned visual gamma (60-90Hz; WOI=[250ms, 500ms]; baseline WOI=[-500ms, -250ms]), instruction cue-aligned visual gamma (60-90Hz; WOI=[100ms, 500ms]; baseline WOI=[-500ms, -100ms]), RDK-aligned sensorimotor beta (15-30Hz; WOI=[0s, 2s]; baseline WOI=[-500ms, 0ms]), response-aligned sensorimotor beta (15-30Hz; WOI=[500ms, 1s]; baseline WOI=[-250ms 250ms]), response-aligned sensorimotor gamma (60-90Hz; WOI=[-100ms, 200ms]; baseline WOI=[-1.5s, -1s]). For each signal, we defined an ROI by comparing power in the associated frequency band during the WOI with a prior baseline WOI at each vertex and averaging over trials. Vertices in either surface with a mean value in the 80^th^ percentile over all vertices in that surface, as well as the corresponding vertices in the other surface, were included in the ROI. This ensured that the contrast used to define the ROI was orthogonal to the subsequent pial versus white matter surface contrast. For each trial, ROI values for the pial and white matter surfaces were computed by averaging the absolute value of the change in power compared to baseline in that surface within the ROI. Finally, a paired t-test was used to compare the ROI values from the pial surface with those from the white matter surface over trials (**Figure 3**). This resulted in positive t-statistics when the change in power was greatest on the pial surface, and negative values when the change was greatest on the white matter surface. All t-tests were performed with corrected noise variance estimates in order to attenuate artifactually high significance values (Ridgway et al., 2012).

The control analyses utilized the same procedure, but each introduced some perturbation to the data. The shuffled analysis permuted the lead fields of the forward model prior to source reconstruction in order to destroy any correspondence between the cortical surface geometry and the sensor data. This was repeated 10 times per session, with a different random lead field permutation each time. Each permutation was then used in the laminar analysis for every low and high frequency signal. The co-registration error analysis introduced a rotation (M=10°, SD=2.5°) and translation (M=10mm, SD=2.5mm) in a random direction of the fiducial coil locations prior to source inversion, simulating between-session co-registration error. This was done 10 times per session, with a different random rotation and translation each time. Again, each perturbation was used in the laminar analysis for every low and high frequency signal. The SNR analysis used a random subset of the available trials from each subject, gradually increasing the number of trials used from 10 to the number of trials available. This was repeated 10 times, using a different random subset of trials each time, and the resulting t-statistics were averaged.

## Condition Comparison

For each visual and sensorimotor frequency band/task epoch combination, induced activity was compared between task conditions on the surface and within the anatomically constrained ROI identified from the corresponding laminar analysis. Seven-cycle Morlet wavelets were used to compute power within the frequency band and this was baseline-corrected in a frequency-specific manner using robust averaging. For each participant, the mean percent change in power over the WOI was averaged over all trials within every condition. Wilcoxon tests for comparing two repeated measures were used to compare the change in power for instruction cue-aligned visual gamma and sensorimotor beta rebound between congruent and incongruent trials. A Friedman test for comparing multiple levels of a single factor with repeated measures was used to compare the sensorimotor beta decrease between low, medium, and high RDK coherence trials. This was followed up by Tukey-Kramer corrected pairwise comparisons. Only trials in which a correct response was made were analyzed.

## Acknowledgements

JB is funded by a BBSRC research grant (BB/M009645/1). SM is supported by a Medical Research Council and Engineering and Physical Sciences Research Council grant MR/K6010/86010/1, the Medical Research Council UKMEG Partnership grant MR/K005464/1, and a Wellcome Principal Research Fellowship to Neil Burgess. SL is supported by a Wellcome Trust clinical postdoctoral grant (105804/Z/14/Z). The WCHN is supported by a strategic award from Wellcome (091593/Z/10/Z).

